# Merlin restoration prevents schwannoma progression in a genetically engineered mouse model of *NF2*-SWN

**DOI:** 10.64898/2026.09.12.751200

**Authors:** Jeremie Vitte, Christine Chiasson-MacKenzie, Banumathi Cole, Andrea I McClatchey, Marco Giovannini

**Author notes:** Corresponding authors: J.V. and M.G.

## Abstract

**Purpose:** Patients with *NF2*-related schwannomatosis (*NF2*-SWN) present with hallmark bilateral vestibular schwannomas, but also schwannomas on other cranial, spinal, and peripheral nerves, as well as meningiomas and ependymomas, caused by germline mutations in the tumor suppressor gene *NF2*. Current therapies involving surgery and radiosurgery are effective for individual tumors but are not always a viable option for patients with multiple tumors, and harbor significant risk of neurological deficits and morbidity. Gene replacement therapy is becoming a promising new treatment strategy for several neurologic diseases. This study aims to understand if restoration of a functional merlin protein, gene product of the *Nf2* gene, in *Nf2*-deficient tumor cells, can provide preclinical therapeutic efficacy in a *NF2*-SWN genetically engineered mouse model.

**Methods:** We have developed a new *Nf2* allele (*Nf2*^*FRT*^) that allows to conditionally restore *Nf2* expression by activation with the Flp recombinase. We generated a new mouse model using the *Nf2*^*FRT*^ allele in combination with *Nf2*^*flox*^ and *Postn-Cre* alleles.

**Results:** Using the new conditional schwannoma mouse model *Postn-Cre;Nf2*^*FRT/flox*^*;R26*^*FlpoER*^, we validated the hypothesis that restoration of *Nf2* reduces the growth of schwannoma.

**Conclusion:** Demonstrating that merlin restoration effectively controls schwannoma growth is a crucial first step toward developing this concept as a new therapeutic strategy for potentially treating schwannomas. The new mouse model will be used to better understand the cellular and molecular mechanisms involved in stopping the growth of *Nf2*-deficient tumors.

## Introduction

Schwannomas and meningiomas are hallmark tumors of *NF2*-related schwannomatosis (*NF2*-SWN), an autosomal dominant tumor predisposition syndrome caused by germline pathogenic variants in the *NF2* tumor suppressor gene and characterized by the development of multiple tumors of the central and peripheral nervous systems, including the hallmarks bilateral vestibular schwannomas (Evans, 2009; Plotkin et al., 2022). Although predominantly histologically benign, these tumors can cause substantial morbidity through compression of cranial, spinal, and peripheral nerves, resulting in progressive hearing loss, neurological deficits, pain, and disability (Ardern-Holmes et al., 2017). *NF2* encodes merlin, a membrane–cytoskeleton-associated tumor suppressor that coordinates cell polarity, contact-dependent growth control, receptor trafficking, and multiple mitogenic signaling pathways (Lallemand et al., 2009; Petrilli and Fernández-Valle, 2016). *NF2* inactivation is also a defining molecular event in the vast majority of sporadic schwannomas and in a substantial subset of sporadic meningiomas (Jacoby et al., 1996). The central role of *NF2* in schwannoma development is further illustrated by non-*NF2* schwannomatosis syndromes associated with germline pathogenic variants in *SMARCB1* or *LZTR1*, in which schwannoma formation frequently involves subsequent somatic biallelic inactivation of *NF2* (Hadfield et al., 2008; Sestini et al., 2008; Kehrer-Sawatzki et al., 2017; Vitte et al., 2017). Thus, despite distinct genetic contexts, loss of merlin represents a convergent and potent driver of schwannoma tumorigenesis.

With only few targeted therapies available (i.e. bevacizumab), current management of schwannomas relies primarily on longitudinal surveillance. Surgical resection is often necessary but carries substantial risk, particularly in patients with *NF2*-SWN who develop multiple, recurrent, or anatomically challenging tumors (Goldbrunner et al., 2020; Rahman et al., 2026; Zalaquett et al., 2026). The development of effective systemic therapies has been complicated in part by the remarkable biological heterogeneity of schwannomas. Despite their relative genetic simplicity and shared dependence on *NF2* inactivation, recent single-cell and histological analyses of human tissues and mouse model have revealed that schwannomas display substantial intratumoral cellular and transcriptional diversity, comprising heterogeneous neoplastic Schwann-cell states, destabilization of Schwann-cell polarity as well as complex stromal and immune populations, thus explaining highly variable growth trajectories, clinical manifestations, histologic features, and therapeutic responses (Chiasson-MacKenzie et al., 2023; Gonzalez Castro et al., 2025; Zhao et al., 2025).

Since acquisition of diverse cellular states and signaling dependencies certainly limit the efficacy of therapies directed against individual downstream pathways, gene-targeted therapy may provide an alternative strategy by intervening at the level of the causal genetic lesion rather than its heterogeneous downstream consequences. Such approaches can restore, modify, replace, or silence a disease-causing gene or its product through technologies including viral and nonviral gene delivery, antisense oligonucleotides, RNA interference, and genome editing (Staedtke et al., 2024; O’Donohue et al., 2025). Gene replacement/restoration has shown clinical success for monogenic disorders, for example SMA (Mendell et al., 2017), making it a particularly attractive therapeutic approach for genetic tumor-predisposition syndromes. For example, multiple proof-of-concept studies with gene-directed approaches have been explored preclinically for the neurofibromatoses, including gene replacement strategies for *NF1* and exon-skipping, cytotoxic or “suicide” gene delivery, and gene replacement approaches for *NF2*-SWN (Yuan et al., 2024; O’Donohue et al., 2025). Most notably, intratumoral administration of an AAV1 vector encoding merlin reduced proliferation and induced regression of human *NF2*-null Schwann-cell-derived xenografts, establishing the therapeutic potential of merlin re-expression in established schwannoma tissue (Prabhakar et al., 2022). Collectively, these findings raise the possibility that restoration of the tumor suppressor gene *NF2* could overcome the complexity and heterogeneity of downstream activated pathways in schwannomas.

However, a fundamental biological question underlying this therapeutic strategy remains unresolved. Although *NF2* inactivation is sufficient to initiate schwannoma formation, whether restoration of merlin could reverse or alter the progression of autochthonous tumors that have adapted to the loss of merlin *in vivo*, and thus have acquired cellular and molecular heterogeneity, specific microenvironmental interactions or adaptive signaling circuits allowing them to escape traditional pathway-targeted strategies, is not known. Here, we developed a genetically engineered mouse model that uses Flp-FRT recombination to enable spatiotemporal restoration of endogenous *Nf2* expression *in vivo*. By combining this system with an established genetically engineered models of schwannoma, we directly tested whether restoration of merlin is sufficient to inhibit the progression of schwannomas. We demonstrate that genetic restoration of *Nf2* impairs schwannoma progression *in vivo* and provide a genetic foundation for the development of *NF2*-restoration strategies for *NF2*-driven tumors.

## Results

### Generation of a new conditional genetically engineered mouse model with tamoxifen-inducible merlin re-expression

To determine whether restoring merlin expression impacts tumor development and possibly induces schwannoma cells stasis or regression *in vivo*, we created a new conditional genetically engineered mouse model (*Postn-Cre;Nf2*^*FRT/flox*^*;R26*^*FlpoER*^) using the *Postn-Cre* (Lindsley et al., 2007) and *Nf2*^*flox*^ (Giovannini et al., 2000) alleles to induce schwannoma development (Gehlhausen et al., 2015; Chiasson-MacKenzie et al., 2023). This mouse model is also carrying a *R26*^*FlpoER*^ allele, expressing a tamoxifen-inducible FlpoER recombinase (Lao et al., 2012) and a new *Nf2*^*FRT*^ allele, allowing re-expression of the endogenous *Nf2* allele after FlpoER recombination. In these mice, treatment with tamoxifen will activate the FlpoER recombinase, which will remove the FRT-STOP-FRT cassette that was blocking the expression of the *Nf2*^*FRT*^ allele, thus allowing expression of *Nf2* from that allele (Figure 1a).

**Figure 1.**
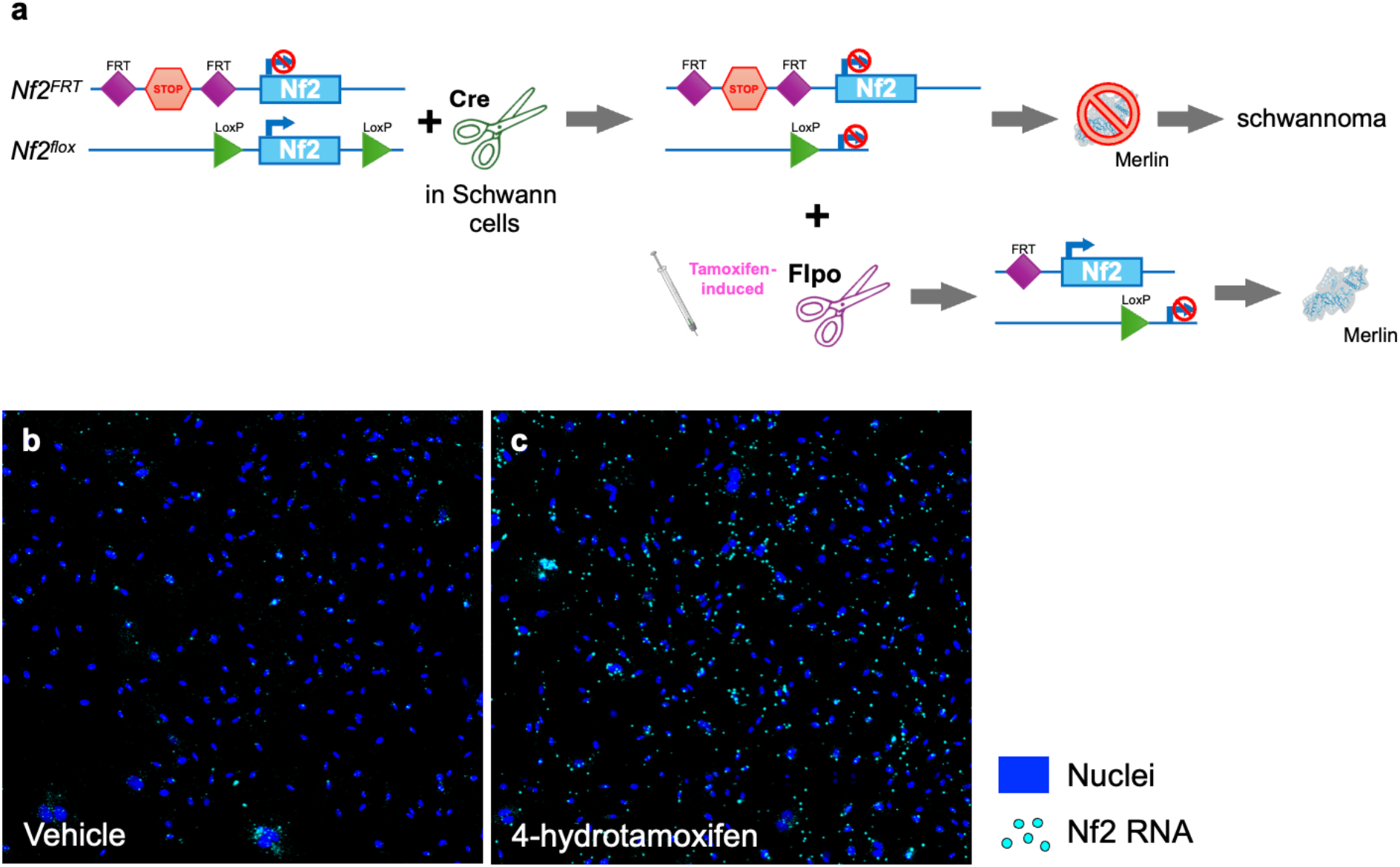
Generation of a new mouse model and *in vitro* validation of *Nf2* re-expression. **a**. Functional schematic of the new conditional genetically engineered mouse model (*Postn-Cre;Nf2*^*FRT/flox*^*;R26*^*FlpoER*^) generating schwannomas by induction of merlin loss in Schwann cells and possibility of merlin re-expression after tamoxifen treatment. **b and c**. *In vitro* validation of *Nf2* re-expression in a primary Schwann cell culture from *Postn-Cre;Nf2*^*FRT/flox*^*;R26*^*FlpoER*^ DRG. The BaseScope assay shows an increased number of cells expressing *Nf2* RNA (blue dots) after treatment with 4-hydroxytamoxifen (c) compared to vehicle (b).

### In vitro validation

To validate the re-expression of *Nf2* in the transgene system described above, we prepared primary Schwann cell cultures isolated from *Postn-Cre;Nf2*^*FRT/flox*^*;R26*^*FlpoER*^ Dorsal Root Ganglia (DRG). The cells were treated with 4-hydroxytamoxifen or vehicle and then stained with a custom designed BaseScope assay targeting the exon 2 of *Nf2* RNA (Figure 1b and c). Overall, the 4-hydroxytamoxifen treatment increased the number of cells expressing Nf2 RNA, thus validating the induction of *Nf2* expression in Schwann cells by 4-hydroxytamoxifen treatment.

### Effect of *Nf2* re-expression in the *Postn-Cre;Nf2*^*FRT/flox*^*;R26*^*FlpoER*^ mouse model

To test the hypothesis that restoration of merlin would impact the development of schwannoma in this model, we generated 2 groups of 6 mice with the following genotypes: *Postn-Cre;Nf2*^*FRT/flox*^*;R26*^*FlpoER*^ and their controls *Nf2*^*FRT/flox*^*;R26*^*FlpoER*^. Mice from each group were treated with tamoxifen (n=3) or vehicle (n=3) at 1 month of age, when lesion is known to start forming with a small but significant increase of cell number in *Postn-Cre;Nf2*^*flox/flox*^ compared to controls (Wright et al., 2026). DRG were dissected from all mice at 6 months of age, when *Postn-Cre;Nf2*^*flox/flox*^ mice have established and well characterized schwannomas with increased tumor cell number, including macrophages and a strongly disrupted architecture with increased intersoma distance (Wright et al., 2026). Similar to *Postn-Cre;Nf2*^*flox/flox*^ mice, *Postn-Cre;Nf2*^*FRT/flox*^*;R26*^*FlpoER*^ mice developed schwannoma lesions with clear increased tumor cell number and presence of whorls (red dashed lines) compared to *Nf2*^*FRT/flox*^*;R26*^*FlpoER*^ control mice (Figure 2). Tamoxifen treatment did not alter the morphology of the DRG of the *Nf2*^*FRT/flox*^*;R26*^*FlpoER/FLTG*^ control mice compared the vehicle-treated group, showing that tamoxifen has no side effect. However, DRG from the tamoxifen-treated *Postn-Cre;Nf2*^*FRT/flox*^*;R26*^*FlpoER*^ group showed strong morphological differences compared to the vehicle-treated *Postn-Cre;Nf2*^*FRT/flox*^*;R26*^*FlpoER*^ group. The tamoxifen-treated DRG showed only a mild increased cell number compared to tamoxifen or vehicle-treated *Nf2*^*FRT/flox*^*;R26*^*FlpoER/FLTG*^ control mice and did not present any whorls (Figure 2).

**Figure 2.**
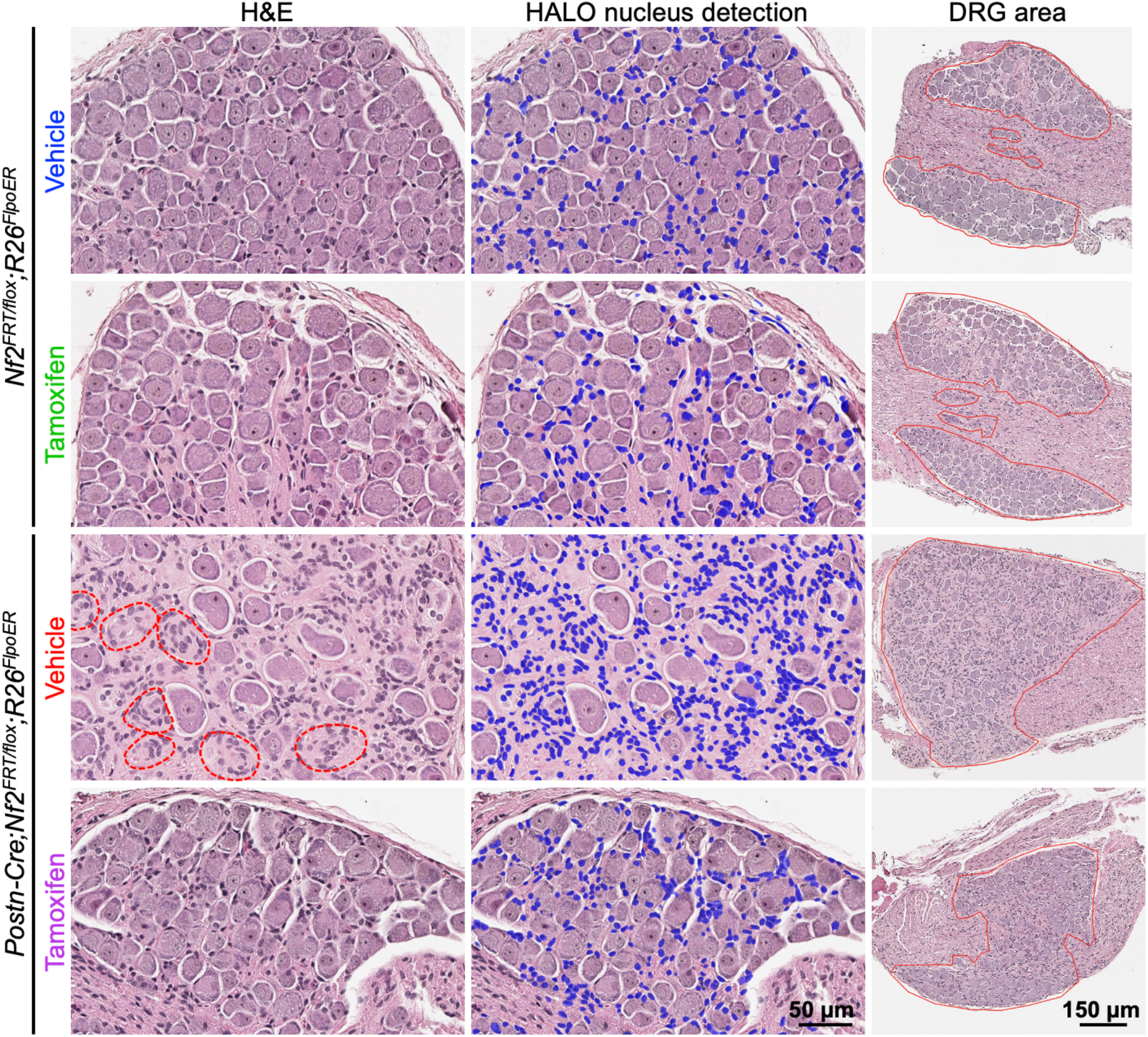
Morphometric analysis of DRG shows that merlin restoration prevents schwannoma growth. H&E staining of DRG from *Nf2*^*FRT/flox*^*;R26*^*FlpoER*^ control and *Postn-Cre;Nf2*^*FRT/flox*^*;R26*^*FlpoER*^ mutant mice treated with vehicle or tamoxifen (1^st^ column), with detection of nuclei by HALO within the neuron cell cluster or within the tumor (blue dots, 2^nd^ column) and DRG/tumor area calculated from manual annotations (red line, 3^rd^ column). Whorls of tumor cells are highlighted with red dashed lines.

To quantify the effect of merlin restoration on the morphology of the DRG schwannomas, we created an algorithm pipeline using the AI-based image analysis HALO software (Indica Labs, Inc). Nuclei (blue dots, Figure 2) were automatically counted in the DRG structures annotated by the user as neuron cell clusters and tumor areas (red line, Figure 2), without counting neuron nuclei. Results confirm the increased DRG cell density and size in the vehicle-treated *Postn-Cre;Nf2*^*FRT/flox*^*;R26*^*FlpoER*^ group, due to the development of the schwannomas, compared to the control groups but also show a significantly decreased tumor cell density and size in the tamoxifen-treated vs vehicle-treated *Postn-Cre;Nf2*^*FRT/flox*^*;R26*^*FlpoER*^ DRG (Figure 3a and b). Additionally, except one outlier, the size of DRG in the tamoxifen-treated *Postn-Cre;Nf2*^*FRT/flox*^*;R26*^*FlpoER*^ group stayed the same as the two control groups (Figure 3b). Overall, these results demonstrate that re-expression of *Nf2* at 1 month of age prevented the growth of the schwannomas.

**Figure 3.**
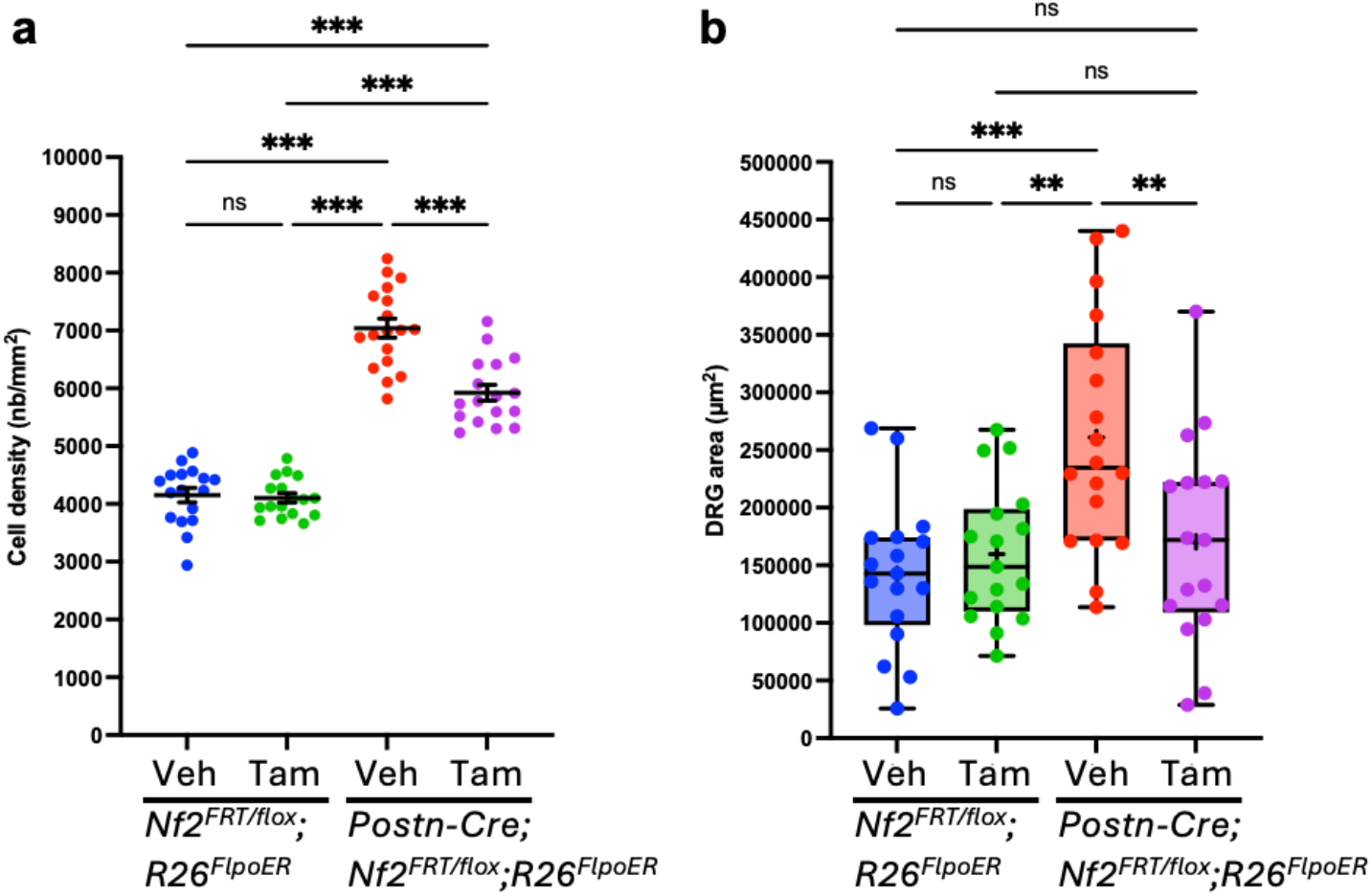
Quantification shows schwannoma growth prevention in the DRG with merlin restoration. Mean tumor cell density (±SEM) (**a**) and DRG area (whiskers showing the minimum to maximum, “+” sign representing the mean) (**b**) for the four experimental groups (n = 3 mice, n = 17-18 DRG per group). Each dot represents a single DRG.

## Discussion

A major strength of the model developed here is the ability to restore endogenous *Nf2* expression with spatial and temporal control. In the current study, we used a ubiquitously expressed FlpoER recombinase, but we could also use a cell-specific promotor to understand the role of a specific cell population. Although this genetic system is not directly translatable to patients, it provides a useful platform to address fundamental questions relevant to the development of *NF2* gene-restoration therapies. In particular, the timing of merlin restoration can be experimentally controlled to determine whether restoration before tumor onset prevents schwannoma formation or, conversely, whether restoration after tumor establishment arrests tumor growth or induces regression. This distinction is important because, although loss of *Nf2* is sufficient to initiate schwannoma formation, it remains unclear to what extent established tumors remain dependent on sustained merlin loss after acquiring additional cellular and molecular changes.

An important feature of this model is that merlin is restored from the endogenous *Nf2* locus rather than expressed from an exogenous construct. Consequently, *Nf2* remains under its endogenous transcriptional regulation, and merlin is restored without supraphysiologic overexpression. This differs from previous gene-replacement studies in which heigh levels of merlin were delivered using AAV viral vectors driven by heterologous promoters in a xenograft model (Prabhakar et al., 2022). Ectopic expression of wild-type merlin in human schwannoma cells has been shown to inhibit proliferation, induce cell-cycle arrest, and promote apoptosis (Schulze et al., 2002; Fraenzer et al., 2003). These findings support the therapeutic potential of merlin restoration but also underscore the importance of controlling the level and duration of transgene expression. Restoration of endogenous *Nf2* in our model provides an opportunity to define the biological consequences of physiologic merlin restoration independently of effects associated with ectopic or supraphysiologic expression. From a therapeutic perspective, these findings suggest that future *NF2* gene-restoration strategies may benefit from approaches that reproduce, as closely as possible, the endogenous regulation and expression levels of merlin.

Our findings further indicate that re-expression of *Nf2* from a single functional allele is sufficient to substantially restrict schwannoma growth. This observation is particularly relevant to gene-replacement strategies, as complete restoration of merlin expression to levels produced by two functional alleles may not be necessary to achieve a therapeutic effect. Determining the minimum level of merlin required for tumor control will nevertheless be important, particularly for gene-delivery approaches in which transduction efficiency and expression levels are likely to vary among tumor cells.

Despite the marked reduction in DRG enlargement following merlin restoration, treated DRG retained a higher cellular density than those of control mice. Thus, merlin restoration strongly restricted tumor growth but did not completely normalize the affected tissue under the conditions examined. The identity of the residual cells remains to be determined. They may represent *Nf2*-deficient tumor cells that escaped recombination and continued to proliferate, *Nf2*-restored tumor cells that persisted despite growth arrest, or non-neoplastic components of the tumor microenvironment. Distinguishing among these possibilities will be important for understanding the cellular response to merlin restoration and the mechanisms underlying incomplete tissue normalization.

Incomplete recombination may also contribute to the residual phenotype. In the current experiments, *Nf2* re-expression was induced using a single course of tamoxifen treatment. Repeated induction could potentially increase the proportion of recombined tumor cells and further suppress tumor growth if a population of neoplastic cells escaped the initial treatment. Quantifying recombination and merlin expression at the single-cell level will therefore be important to distinguish incomplete target engagement from biological resistance to *Nf2* re-expression. Since heterogeneous delivery to tumor cells will represents an important challenge in patients, understanding the minimal fraction of tumor cells that must be transduced to control the entire tumor progression or induce regression is directly relevant to the translation of gene therapy to *NF2*-SWN patients.

An important next step will be to determine the consequences of *Nf2* re-expression after tumors are fully established. The experiments presented here primarily demonstrate that merlin restoration restricts schwannoma progression; however, treatment of established tumors will more closely model the clinical setting in which gene-restoration therapy would be administered. Longitudinal studies in which merlin is restored at different stages of tumor development should determine whether the response depends on tumor age, size, or degree of cellular heterogeneity and, critically, whether restoration arrests further growth or causes regression of an existing tumor.

The cellular mechanism underlying the response to *Nf2* re-expression also remains to be established. Merlin restoration could induce a durable proliferative arrest, promote differentiation or senescence, or lead to elimination of tumor cells through apoptosis. Previous studies in human schwannoma cells have shown that ectopic expression of wild-type merlin reduces proliferation, promotes G0/G1 arrest, and increases apoptosis, while AAV-mediated merlin replacement in *NF2*-null schwannoma xenografts was associated with reduced proliferation and increased tumor-cell apoptosis (Schulze et al., 2002; Fraenzer et al., 2003; Prabhakar et al., 2022). Determining whether endogenous *Nf2* re-expression produces similar responses in autochthonous schwannomas will be important for understanding the durability of tumor control. A predominantly cytostatic response could require sustained merlin expression to prevent renewed tumor growth, whereas elimination of neoplastic cells through apoptosis could potentially produce a more durable response. These distinctions have important implications for the design, dosing, and duration of future *NF2* gene-restoration therapies.

## Material and methods

### Mice, generation of the *Nf2*^*FRT*^ allele and tamoxifen treatment

All mouse strains were maintained on FVB/N genetic background. All animal care and experimentation were performed with the approval of the UCLA Institutional Animal Care and Use Committees under protocol number 2019-018. Mice were housed under standard conditions: 12 hours of light/12 hours of dark; ambient temperature range 20–26 °C; ambient humidity range 30–70%. Mice were monitored twice a week until dissection at 6 months of age. Mice were euthanized by CO2 inhalation. The *Nf2*^*FRT*^ allele was constructed by inserting a “FRT-STOP-FRT” cassette (FRT-PGK-Puro-SV40PolA-FRT) in intron 1 of the *Nf2* gene, using BamHI restriction sites. Mice were genotyped by TransnetYX (Cordova, TN) or in house by PCR amplification of DNA extracted from tail biopsies using the primers cre-s1 (5′-ACA TGT TCA GGG ATC GCC AG-3′) and cre-a1 (5′-TAA CCA GTG AAA CAG CAT TGC-3′) for the *Cre* transgene, NF2Flox2-S (5′-CTT CCC AGA CAA GCA GGG TTC-3′) and NF2Flox2-A (5′-GAA GGC AGC TTC CTT AAG TC-3′) for the *Nf2*^*flox*^ and Nf2^WT^ alleles, Nf2-FRT-F (5’-CACCTGCTCCAGCACACA-3’) and Nf2-FRT-R (5’-CAGAAGCTGGTCGAGGAAGTT-3’) for the *Nf2*^*FRT*^ allele, 10507 (5’-TTATGTAACGCGGAACTCCA-3’), oIMR8545 (5’-AAAGTCGCTCTGAGTTGTTAT-3’) and oIMR8546 (5’-GGAGCGGGAGAAATGGATATG-3’) for the *R26*^*FlpoER*^ and *R26*^*WT*^ alleles. Mice were treated at 1 month of age for 5 consecutive days with 2 mg of tamoxifen (Sigma T-5648) resuspended in corn oil (Sigma C-8267) or an equivalent volume of corn oil for the vehicle group, by intraperitoneal injection.

### Primary cell culture, 4-Hydroxytamoxifen treatment and BaseScope Assay

Primary Schwann cell cultures were isolated from cervical and lumbar DRG collected from a *Postn-Cre;Nf2*^*FRT/flox*^*;R26*^*FlpoER*^ mouse at 8 months of age, dissociated with collagenase type I (Invitrogen 17100-017) and dispase II (Roche 04 942 078 001), and plated in Poly-L-Lysin (Sigma P4707)/laminin (Invitrogen 23017-015)-coated cell culture flasks in PM medium (DMEM, FBS, amphotericin B, forskolin, neuregulin and N2 supplement). Cells were then passaged and seeded into chambered coverglass slides (Lab-Tek cat#155411), treated with 4-Hydroxytamoxifen (Sigma H7904) or vehicle (Ethanol 100%) for 2 days. Slides were processed for the BaseScope v2 RED Assay (Bio-Techne 323900) following manufacturer instructions and using a custom probe targeting mouse *Nf2* exon 2. Images were taken by confocal microscopy using the fluorescence signal emitted by the Fast Red precipitate.

### H&E staining and cell counting

Cervical DRG were dissected, fixed with 10% neutral buffered formalin for 24 hours and processed using standard protocols into Formalin-Fixed Paraffin-Embedded (FFPE) blocks, arranged in a “DRG array”. For each block, about 40-50 3.5µm-thick sections were collected and only the 10 sections containing the higher surface of DRG tissues were kept for further processing. H&E staining was performed following standard procedures. Whole H&E slides were scanned by the UCLA TPCL core facility using a Leica Aperio scanner. QuPath v0.6.0 (Bankhead et al., 2017) was used for DRG annotations and HALO AI v4.0.5107 (Indica Labs, Albuquerque, NM) was used with customized algorithms to recognize the neurons and count only non-neuronal cells in the annotated tissue layers.

### Statistics

Raw data from HALO was processed using standard core packages in R (R Core Team, 2026) and imported to Prism v11.0.0 (GraphPad Software, Boston, MA). Unpaired two tailed Welch’s t test was used to compare differences between two groups.

## Acknowledgements

We would like to thank Dr. Fausto J. Rodriguez (UCLA, Department of Pathology) for histopathological review of mouse schwannomas and Gabrielle Thompson for technical support. Confocal laser scanning microscopy was performed at the Advanced Light Microscopy/Spectroscopy Laboratory (RRID: SCR_022789) and the Leica Microsystems Center of Excellence at the California NanoSystems Institute at UCLA with funding support from NIH Shared Instrumentation Grant S10OD025017 and NSF Major Research Instrumentation grant CHE-0722519. This work was supported by the U.S. Army Medical Research and Development Command, through the Neurofibromatosis Research Program under Award Nos. W81XWH-21-1-0446 (A.I.M.) and W81XWH-21-1-0448 (M.G.). Opinions, interpretations, conclusions, and recommendations are those of the authors and are not necessarily endorsed by the Department of Defense.

